# Re-evaluating the salty divide: phylogenetic specificity of transitions between marine and freshwater systems

**DOI:** 10.1101/347021

**Authors:** Sara F. Paver, Daniel J. Muratore, Ryan J. Newton, Maureen L. Coleman

## Abstract

Marine and freshwater microbial communities are phylogenetically distinct and transitions between habitat types are thought to be infrequent. We compared the phylogenetic diversity of marine and freshwater microorganisms and identified specific lineages exhibiting notably high or low similarity between marine and freshwater ecosystems using a meta-analysis of 16S rRNA gene tag-sequencing datasets. As expected, marine and freshwater microbial communities differed in the relative abundance of major phyla and contained habitat-specific lineages; at the same time, however, many shared taxa were observed in both environments. *Betaproteobacteria* and *Alphaproteobacteria* sequences had the highest similarity between marine and freshwater sample pairs. *Gammaproteobacteria* and *Alphaproteobacteria* contained the highest number of Minimum Entropy Decomposition nodes shared by marine and freshwater samples. Shared nodes included lineages of the abundant alphaproteobacterial group SAR11 that have not previously been reported in 16S rRNA gene surveys of freshwater lakes. Our results suggest that shared taxa are numerous, but tend to occur sporadically and at low relative abundance in one habitat type, leading to an underestimation of transition frequency between marine and freshwater habitats. Lineages with a high degree of shared taxa or habitat-specific diversification represent targets for genome-scale investigations into microbial adaptations and evolutionary innovations. Rare taxa with abundances near or below detection, including lineages that appear to have crossed the salty divide relatively recently, may have novel adaptations enabling them to exploit opportunities for niche expansion when environments are disturbed or conditions change.

**Importance:** The distribution of microbial diversity across environments yields insight into processes that create and maintain this diversity as well as potential to infer how communities will respond to future environmental changes. We integrated datasets from dozens of freshwater lake and marine samples to compare diversity across open water habitats differing in salinity. Our novel combination of sequence-based approaches revealed phyla and proteobacterial classes inferred to include more or less recent transitions across habitat types as well as specific lineages that are shared by marine and freshwater environments at the level of 16S rRNA sequence types. Our findings contribute to understanding the ecological and evolutionary controls on microbial distributions, and open up new questions regarding the plasticity and adaptability of particular lineages.

## Introduction

Phylogenetic relationships of organisms within and across ecosystems can provide insight into how these organisms evolved and how evolution might proceed into the future. Microorganisms in the water columns of freshwater and marine ecosystems provide a unique juxtaposition. On one hand, these habitats share common features of pelagic lifestyles like free-living and particle-associated niches (1), potential for interactions with phytoplankton (2), and opportunities for diverse photoheterotrophic organisms including aerobic anoxygenic phototrophs (3) and rhodopsin-containing bacteria (4, 5). However, salinity preference is considered a complex trait involving many genes and complex cellular integration (6), suggesting that transitions between high and low salinity are difficult from a genetic perspective. Consistent with this idea, there is evidence for evolutionary separation between microbial lineages in high and low salinity environments and the current paradigm is that transitions between marine and freshwater ecosystems are infrequent, despite their ecological similarities (7).

Environmental sequence data provide support for a salty divide separating marine and freshwater microbial assemblages. Saline environments are compositionally distinct from non-saline environments (8, 9). Salinity-induced shifts in microbial diversity have been observed in studies of marine-to-freshwater gradients in many systems, including the Baltic Sea (10), Columbia River Estuary system (11), and Antarctic lakes that have become progressively less saline since becoming isolated from the sea (12). Moreover, a number of microbial lineages appear to be unique to freshwater lakes (13, 14). For freshwater lineages that are found in multiple habitats, the secondary habitat is most often terrestrial, not marine (14). A review of available studies comparing microorganisms from marine and freshwater environments found evidence for evolutionary separation between marine and freshwater lineages in most studies (7). Observed separation between marine and freshwater lineages and ecosystem-specific sequence clusters form the basis for the conclusion that transitions between marine and freshwater environments are infrequent and most transition events occurred a long time ago (7).

Difficulty detecting transitions between marine and freshwater systems may contribute to the paradigm that transitions occur infrequently. Detecting a transition requires sufficiently abundant extant descendants. Most immigrant cells are expected to go extinct locally due to ecological drift, just as most mutations are lost from a population due to genetic drift (15). The probability of an immigrant avoiding extinction due to ecological drift, like a mutation avoiding genetic drift, depends on the degree of selective advantage. For example, in populations of *E. coli* (~3 × 107 cells; 16), a mutation conferring a 10% advantage appears an average of five times before it is established compared to a mutation with a 0.1% advantage which would need to appear 500 times to avoid extinction by drift (17). In addition to overcoming ecological drift, the degree of selective advantage for cells migrating between marine and freshwater habitats would need to be strong enough to overcome any salinity-based disadvantages. Microorganisms that become established must also achieve sufficiently high population abundances to be reliably detected by current sequencing methods. As amplicon sequencing datasets accumulate from an increased diversity of environments and library size increases, our ability to detect transitions improves.

Here we revisit classic questions concerning divisions between marine and freshwater microorganisms by comparing 16S rRNA V4 region amplicon sequences from available marine and freshwater datasets. This meta-analysis is timely given the accumulation of aquatic 16S rRNA amplicon sequencing datasets and, specifically, the availability of sequence data from the Laurentian Great Lakes. Microbial ecology studies of large lakes, including the Laurentian Great Lakes, have the potential to help bridge gaps in our understanding of marine and freshwater microbes. Large lakes are less influenced by their catchment than smaller lakes and experience oceanic type-physical processes, including strong currents and upwelling (18). Further, extreme oligotrophic conditions in certain locations within the Great Lakes resemble nutrient conditions in some ocean basins (19-21). Our specific objectives were to 1) compare the diversity of marine and freshwater microorganisms and 2) identify lineages that have a comparatively high or low degrees of sequence similarity between marine and freshwater ecosystems. These lineages may represent targets for exploring physiological and molecular barriers to salinity tolerance, and for identifying novel strategies for overcoming these barriers.

## Results

### Marine and freshwater taxa are distinct at varying phylogenetic resolution

Marine microbial assemblages were phylogenetically distinct from freshwater assemblages based on UniFrac comparisons of MED nodes from marine and freshwater samples (Fig. S1; Weighted Unifrac PerMANOVA F=23.6, R^2^=0.24, p<0.001, df=75; Unweighted Unifrac PerMANOVA F=41.8, R^2^=0.36, p<0.001, df=75). Phyla (classes for *Proteobacteria*) including *Alphaproteobacteria*, *Gammaproteobacteria*, *Euryarchaeota*, and *Marinimicrobia* had significantly higher relative abundances in marine systems while *Betaproteobacteria* and *Verrucomicrobia* had higher relative abundances in freshwater systems (Fig. 1; Table S1). Relative abundance differences between marine and freshwater systems became more pronounced with increasing phylogenetic resolution from phylum to order to family (Fig. S2). The range of unweighted UniFrac distances for pooled marine and freshwater sequences also increased from phylum/ class to family (Fig. S3; Table S1), indicating that phyla and proteobacterial classes tend to include a mix of shared and habitat-specific lineages while certain families contain a higher concentration of shared lineages or a higher concentration of habitat-specific lineages. Gammaproteobacterial families *Chromatiaceae* and *Vibrionaceae* and actinobacterial families PeM15 and *Mycobacteriaceae* had the smallest unweighted UniFrac distances (< 0.55), suggesting that marine and freshwater lineages tend to be more closely related in these groups than other groups. In contrast, *Hydrogenophilaceae* (*Betaproteobacteria*) and KI89A (*Gammaproteobacteria*) each had UniFrac distances of 1.00 (Table S1), indicating that marine and freshwater lineages we detected within these families were completely distinct phylogenetically.

**Figure 1.**
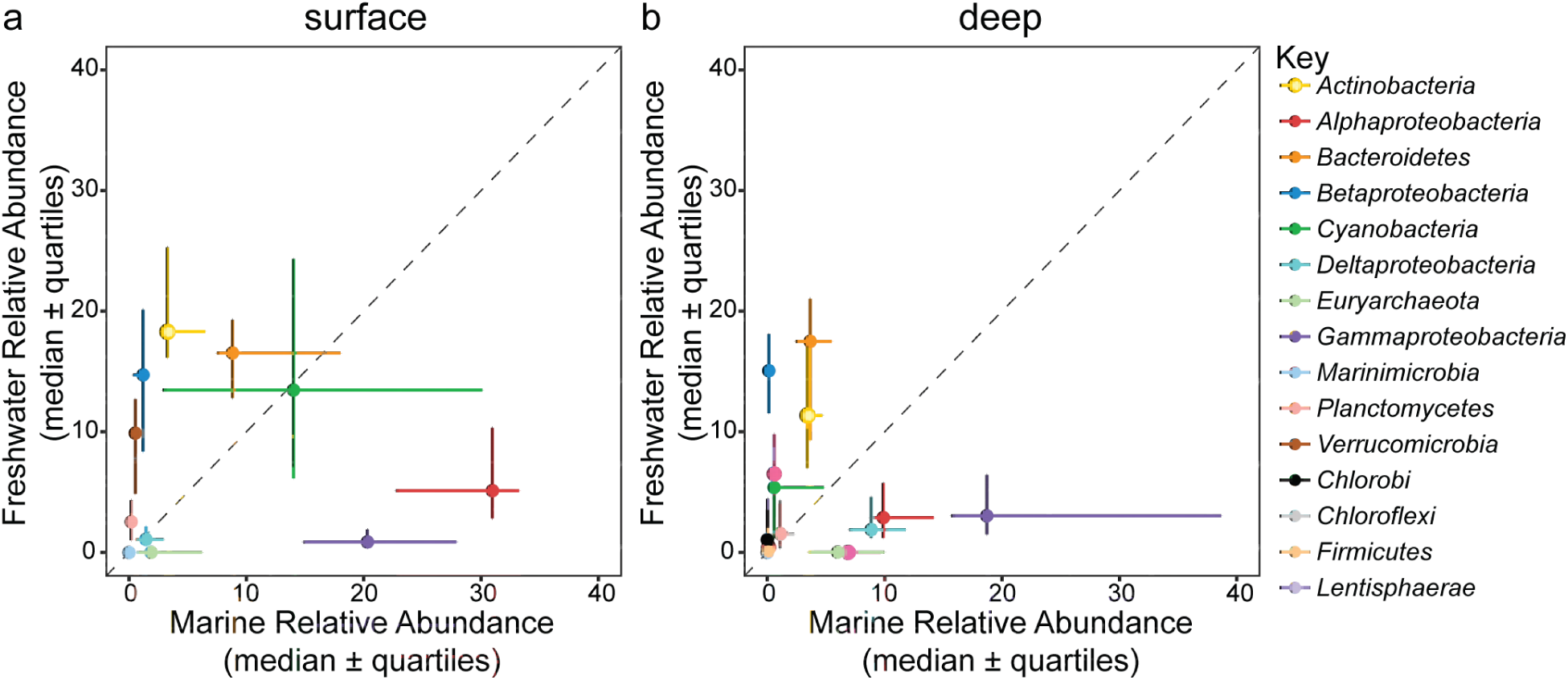
Median relative abundance of phyla and proteobacterial classes in freshwater and marine samples collected from surface (a) and deep (b) waters. The deepest hypolimnion (below thermocline) sample collected from stratified lakes and marine samples collected at depths >75m were classified as ‘deep’ samples. Diagonal lines indicate a 1:1 relationship.

### Direct sequence-level comparisons reveal variation across phyla

To estimate taxon relatedness across habitats independent of operational definitions, we implemented a direct sequence-based comparison to assess the relative phylogenetic distance (as a proxy for time) since the most recent marine-freshwater transition. Within each phylum (class for *Proteobacteria*) we clustered sequences at every possible threshold of sequence identity (0, 1, 2, 3, etc. mismatches) and identified the highest sequence identity threshold at which a shared cluster was detected for each pairwise combination of marine and freshwater samples. For example, if the same exact sequence was found in both the marine and freshwater sample, the threshold would be 1 (100% sequence identity), whereas a deep branching split into exclusively marine and freshwater clades would yield a much lower threshold. This procedure was then repeated for all pairwise combinations of freshwater and marine samples.

Using this approach, we found that most phyla and proteobacterial classes contained shared taxa at identity thresholds >0.99 for at least some pairs of marine and freshwater samples (Fig. 2), suggesting widespread recent transitions across habitat types. We did not detect specific sample pairs preferentially yielding high similarity sequences; sequences with the highest similarity across habitats for each taxonomic group were distributed across the majority of marine and freshwater samples. We found that the median sequence identity threshold for shared taxa across all sample pairs varied strongly across phyla (Fig. 2; Figs. S4 and S5). The majority of marine-freshwater sample pairs contained shared betaproteobacterial and alphaproteobacterial taxonomic units at a comparatively small sequence identity value of 0.96. At the other extreme, no marine-freshwater sample pairs contained shared *Nitrospirae* taxa at identity thresholds >0.89. Moreover, most marine-freshwater sample pairs did not contain shared *Chloroflexi*, *Euryarchaeota*, or *Chlorobi* at identity thresholds >0.77.

**Figure 2.**
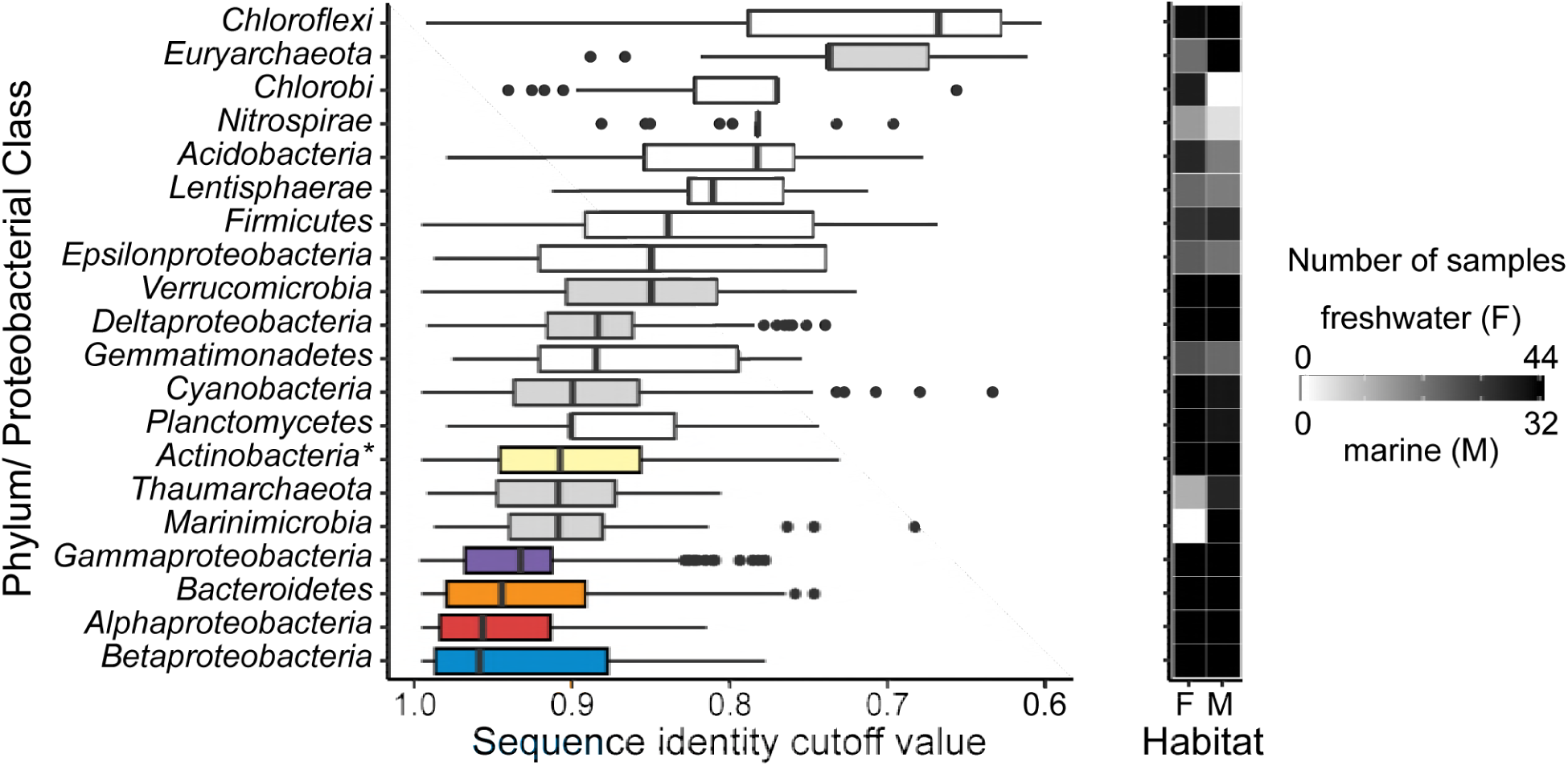
Minimum sequence identity threshold (i.e., finest scale resolution) at which pairs of marine and freshwater samples share common taxa. Box plots indicate the median, quartiles, and range of values observed for all marine and freshwater sample pairs. Colored boxes indicate phyla/ proteobacterial classes that contain 5 or more shared MED nodes while grey boxes indicate groups that contain 1-3 shared MED nodes. A heatmap illustrates the number of freshwater (F) and marine (M) samples containing representatives of each phyla/ proteobacterial class. \**Actinobacteria* cutoff values were calculated with a pre-clustered dataset (see Fig S5 for comparison of all groups using a pre-clustered dataset).

### Abundant phyla consistently contain shared marine-freshwater nodes

We observed 171 total shared MED nodes, i.e. nodes detected in at least one marine sample and one freshwater sample (Fig. 3; Table 1; Fig. S7). For most phyla and proteobacterial classes, a small positive relationship was observed between the number of sample locations included in the analysis and the percent of nodes shared between marine and freshwater samples (Fig. S7). *Betaproteobacteria* and *Gammaproteobacteria* showed the largest increases in shared nodes as more marine and freshwater sites were sampled, respectively. The large increase in shared nodes as more marine sites were sampled suggests that betaproteobacterial diversity was well sampled in freshwater datasets where *Betaproteobacteria* are abundant (Fig. 1) and that betaproteobacterial nodes that are infrequently observed in marine samples originated from freshwater. Similarly, gammaproteobacterial diversity appears to be relatively well sampled in marine datasets where *Gammaproteobacteria* are abundant (Fig. 1) with gammaproteobacterial nodes infrequently observed in freshwater samples likely originating from marine systems. *Gammaproteobacteria* contained the most shared nodes, which accounted for 10% and 33% of total gammaproteobacterial nodes observed in marine and freshwater systems, respectively. *Alphaproteobacteria* contained the second highest number of shared nodes, accounting for 7% and 14% of total alphaproteobacterial nodes observed in marine and freshwater systems, respectively. Notably, shared nodes from marine alphaproteobacterial SAR11 clades surface 1 and surface 2 were observed in a humic lake, an estuarine clade was observed in a Tibetan Plateau lake, and an unclassified SAR11 clade was observed in the Laurentian Great Lakes (Fig. 4).

**Figure 3.**
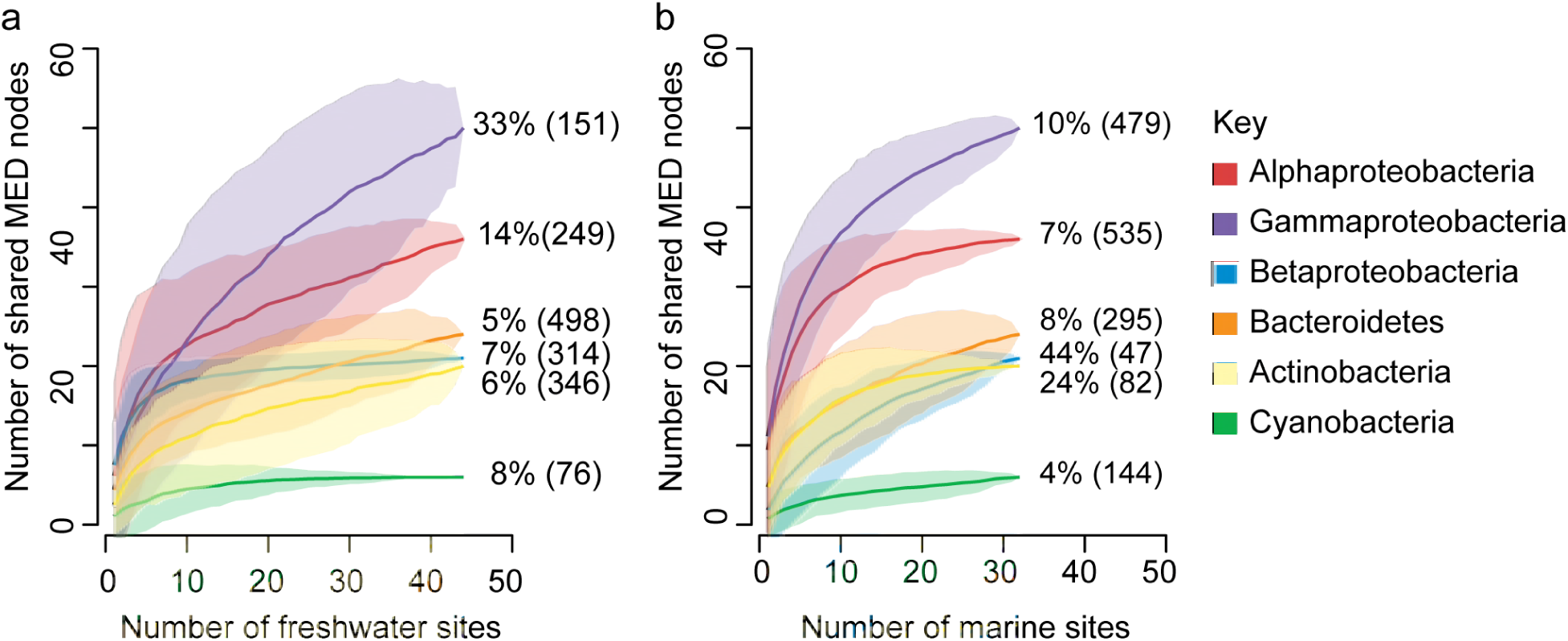
Species accumulation curves for taxonomic groups that contain shared marine and freshwater MED nodes as the number of freshwater sites included in the analysis increases (a) and as the number of marine sites included in the analysis increases (b). The percent of sequences shared between habitats with all sites analyzed is included to the right of each curve; the total number of MED nodes within each group in freshwater and marine habitats, respectively is indicated in parentheses.

**Table 1.** Genera containing at least two shared MED nodes.

| Phylum/ class | Order | Family | Genus | # |
| --- | --- | --- | --- | --- |
| Actinobacteria | Acidimicrobiales | OM1_clade | Ca. Actinomarina | 2 |
| Actinobacteria | Acidimicrobiales | Sva0996_marine_group |  | 2 |
| Actinobacteria | Corynebacteriales | Mycobacteriaceae | *Mycobacterium* | 2 |
| Actinobacteria | Micrococcales | Microbacteriaceae | Ca. Aquiluna | 4 |
| Actinobacteria | PeM15 |  |  | 4 |
| Bacteroidetes | Flavobacteriales | Cryomorphaceae | *Fluviicola* | 5 |
| Bacteroidetes | Sphingobacteriales | Chitinophagaceae | *Sediminibacterium* | 2 |
| Bacteroidetes | Sphingobacteriales | NS11-12_marine_group |  | 3 |
| Cyanobacteria | Subsection I | Family I | *Synechococcus* | 3 |
| Marinimicrobia |  |  |  | 4 |
| Alphaproteobacteria | Caulobacterales | Caulobacteraceae | *Brevundimonas* | 4 |
| Alphaproteobacteria | Rhizobiales | Methylobacteriaceae | *Methylobacterium* | 3 |
| Alphaproteobacteria | Sphingomonadales | Sphingomonadaceae | *Novosphingobium* | 2 |
| Alphaproteobacteria | Sphingomonadales | Sphingomonadaceae | *Sphingobium* | 3 |
| Alphaproteobacteria | Sphingomonadales | Sphingomonadaceae | *Sphingomonas* | 3 |
| Betaproteobacteria | Burkholderiales | Burkholderiaceae | *Ralstonia* | 2 |
| Betaproteobacteria | Burkholderiales | Comamonadaceae | *Aquabacterium* | 2 |
| Deltaproteobacteria | SAR324_clade |  |  | 3 |
| Gammaproteobacteria | Alteromonadales | Alteromonadaceae | *Marinobacter* | 2 |
| Gammaproteobacteria | E01-9C-26_marine_group |  |  | 2 |
| Gammaproteobacteria | Oceanospirillales | Oceanospirillaceae | *Pseudohongiella* | 4 |
| Gammaproteobacteria | Oceanospirillales | OM182_clade |  | 2 |
| Gammaproteobacteria | Oceanospirillales | SAR86_clade |  | 2 |
| Gammaproteobacteria | Pseudomonadales | Moraxellaceae | *Acinetobacter* | 5 |
| Gammaproteobacteria | Pseudomonadales | Pseudomonadaceae | *Pseudomonas* | 3 |
| Gammaproteobacteria | Vibrionales | Vibrionaceae | *Vibrio* | 2 |
| Euryarchaeota | Thermoplasmatales | Marine Group II |  | 2 |
| Thaumarchaeota | Unknown_Order | Unknown_Family | Ca. Nitrosopumilus | 2 |

**Figure 4.**
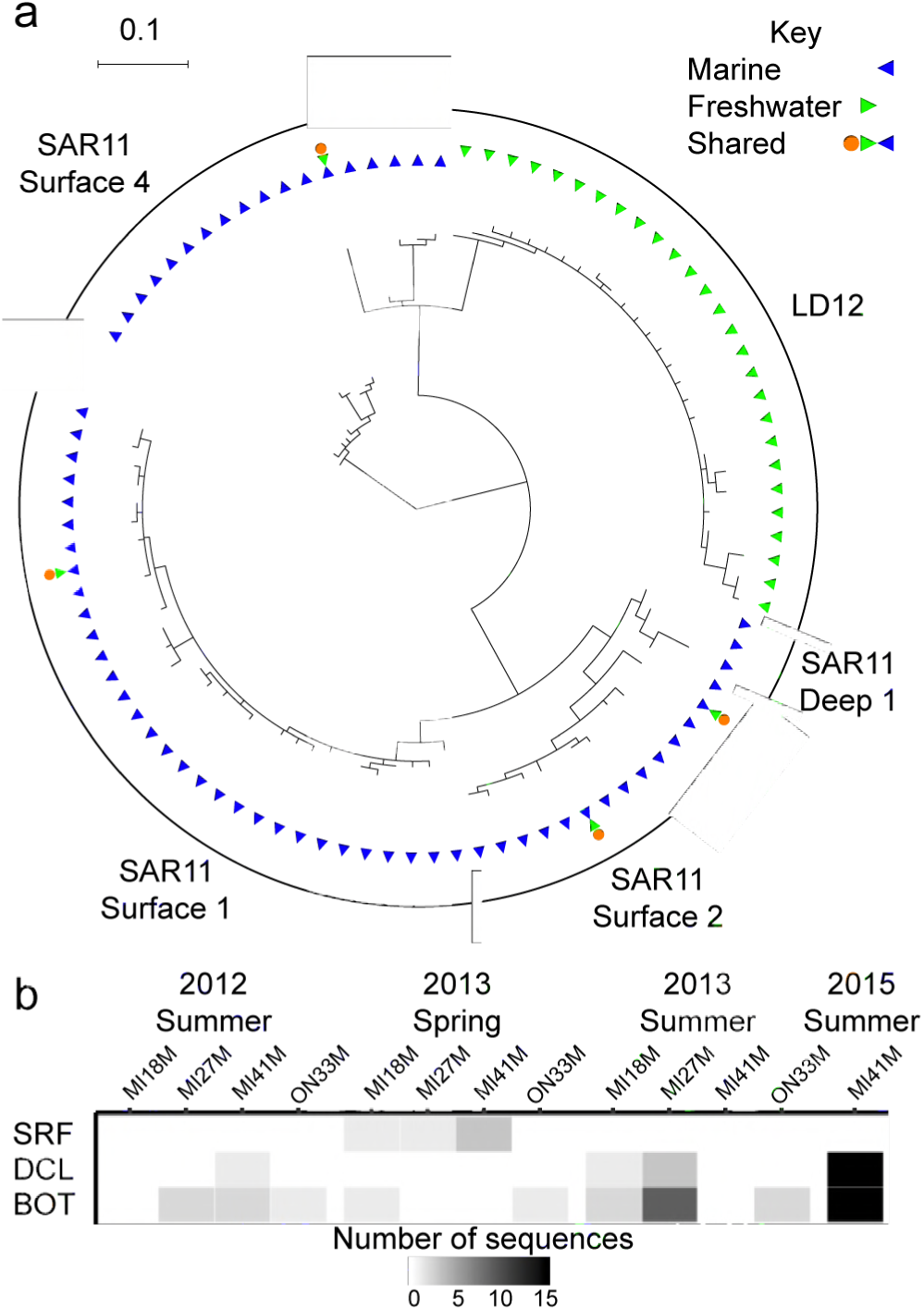
Observations of non-LD12 SAR11. (a) 16S V4 region gene tree constructed using representative sequences from eachSAR11 node. The first ring indicates whether nodes were found only in marine (blue) or freshwater samples (green) while the second ring indicates the nodes that are shared across habitat type (orange). (b) Number of non-LD12 SAR11 clade sequences detected in two stations on Lakes Michigan and Ontario.

### Metagenomic evidence of non-LD12 SAR11 in the Great Lakes

To further investigate SAR11 nodes shared between marine and freshwater habitats, we analyzed the abundance of non-LD12 Pelagibacterales using 16S rRNA data from additional Laurentian Great Lakes samples and surface metagenomes collected from each of the five Great Lakes. For simplicity, we use ‘LD12’ to refer to the common freshwater SAR11 IIIb clade and ‘marine SAR11’ to refer to all other SAR11 clades. When observed in 16S rRNA datasets using the 515F/806R primer set, marine SAR11 sequences accounted for 1-15 sequences out of an average of approximately 75,000 sequences per sample (Fig. 4b). To test whether marine-like sequences could be identified genome-wide, not just at the level of 16S rRNA, we extracted metagenome reads with a top hit of Pelagibacterales from Great Lakes samples and a Tara Oceans sample collected near Bermuda for comparison, assigned them to SAR11 clade protein clusters, and classified them as marine SAR11 or LD12 using pplacer (22). We included all protein clusters with more than 100 sequences identified as marine SAR11 or LD12 at a likelihood value of 0.95 for each metagenome; approximately 300 protein clusters per metagenome satisfied this criterion.

For each metagenome analyzed, 58-59% of sequences within a protein cluster (median value across all protein clusters) were classified as either LD12 or SAR11 at a likelihood of 0.95. Of the classified sequences, 11-12% (median value across all protein clusters) were classified as marine SAR11 in each of the Great Lakes samples, while 98% were classified as marine SAR11 in the marine sample (Fig. 5; Fig. S8). Across protein clusters, the fraction of sequences classified as SAR11 was much more variable for Great Lakes samples (range 0-57%, interquartile range: 6-18%) than for the marine sample (range: 76-100%, interquartile range: 96-99%). We identified 199 protein clusters with more than 5% of sequence reads classified as SAR11 in all five Great Lakes metagenomes, suggesting that similarities between non-LD12 SAR11 in the Great Lakes and marine SAR11 extend beyond the 16S rRNA gene to a substantial portion of the genome.

**Figure 5.**
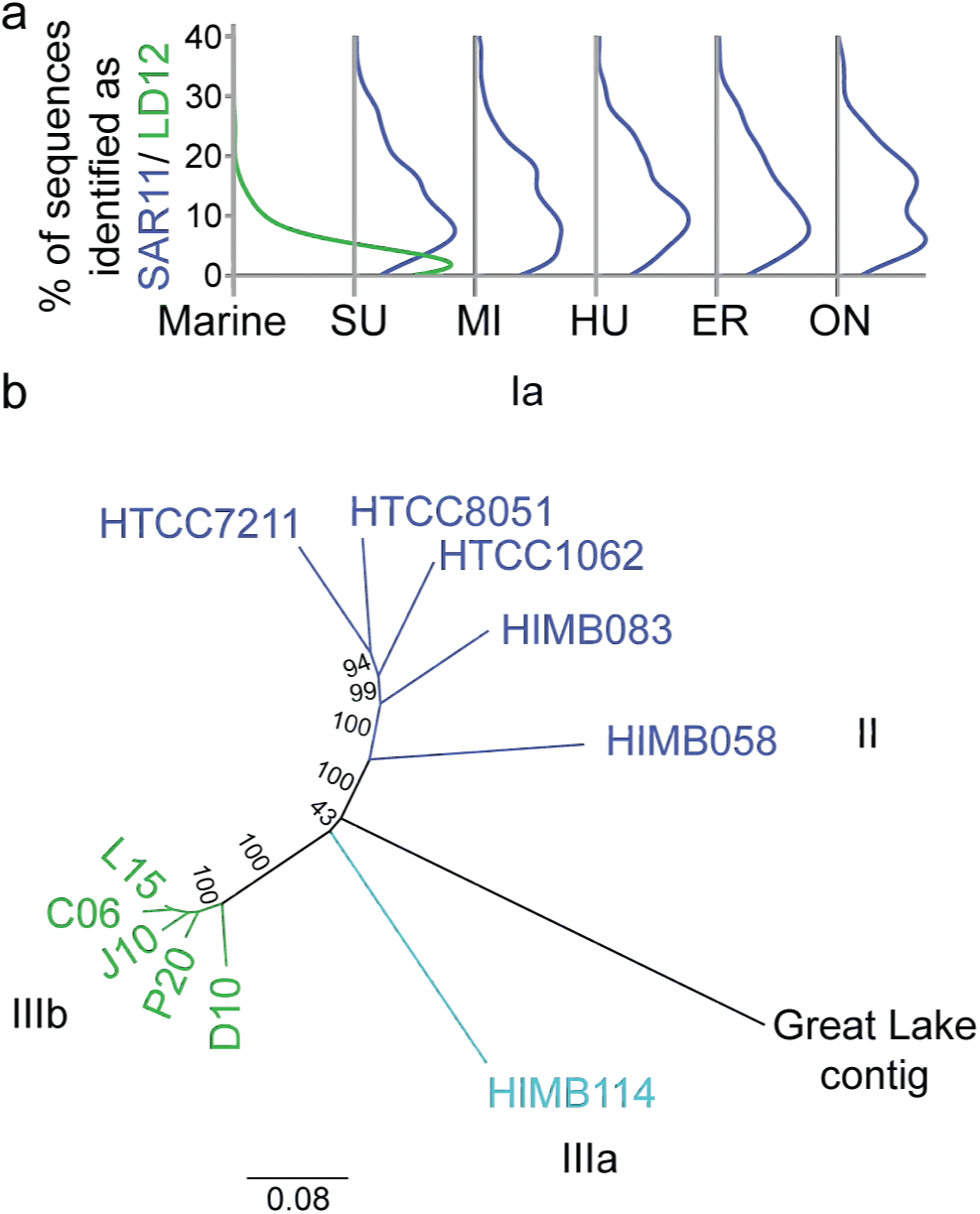
Metagenomic evidence for non-LD12 SAR11 in the Laurentian Great Lakes. (a) Percent of classifed reads identified as LD12 (green) in a Tara Oceans sample collected near Bermuda (marine) and marine SAR11 (blue) in each of the five Laurentian Great Lakes (SU=Superior; MI=Michigan; HU=Huron; ER=Erie; ON=Ontario). Ridge plots present the distribution of identified reads across all protein clusters with greater than 100 reads classified as SAR11 or LD12 at a liklihood value of 0.95. (b) Neighbor-joining consensus tree of 1.2 KB nucleotide sequences from the protein cluster identified as COG2609 (Pyruvate dehydrogenase complex, dehydrogenase E1 component). Strain names are colored based on phylogenetic classification within the SAR11 clade: green - LD12 sequences from group IIIb; light blue - group IIIa, sister group to IIIb; blue - all other marine SAR11 clades included in the analysis (Ia, II); black - a contig assembled from the Lake Erie metagenome. Consensus support values (%) are indicated on branches

## Discussion

We compared marine and freshwater microbial lineages from publicly available 16S rRNA V4 region amplicon sequencing datasets to revisit the paradigm of infrequent transitions between marine and freshwater habitats and identify lineages that may have transitioned between environments with higher or lower frequency than average. At a broad level, marine and freshwater assemblages are phylogenetically distinct. While there is a great deal of overlap in the presence of phyla and proteobacterial classes, some groups are more abundant in one habitat than the other and certain lineages within those groups appear to be habitat-specific. Our results corroborate the findings of Lozupone and Knight (8) and Thompson and colleagues (9) that salinity is the major environmental determinant separating free-living bacteria from different environments. Further, our finding of higher relative abundances of *Betaproteobacteria* and *Actinobacteria* in lakes and higher relative abundances of *Alphaproteobacteria* and *Gammaproteobacteria* in marine systems corresponds with taxonomic comparisons made using metagenomic sequence datasets (23) as well as previous observations using 16S rRNA sequences (14, 24).

We took multiple approaches to assess phylogenetic relationships between marine and freshwater microorganisms within taxonomic groups. We used unweighted UniFrac distances which have been employed previously to make inferences about the frequency of transitions between marine and freshwater systems (7), direct sequence comparisons to generate an estimate of taxon relatedness independent of operational definitions, and comparisons of the number of shared MED nodes with the inference that each shared node encompasses at least one transition between environments. The presence of habitat-specific lineage diversification increases UniFrac distances whether or not a group has a high frequency of shared nodes, making UniFrac distance ill-suited for making inferences about transition frequency (25). Hence we used sequence similarity and number of shared MED nodes to make inferences about taxonomic groups that have the most recent and frequent transition between marine and freshwater.

### Taxa with comparatively high transition frequency

*Alphaproteobacteria*, *Betaproteobacteria*, *Bacteroidetes*, *Gammaproteobacteria* and *Actinobacteria* are the groups we inferred to have the most frequent transitions between marine and freshwater systems. *Alphaproteobacteria* and *Gammaproteobacteria* were two of the most abundant phyla in marine systems while *Actinobacteria*, *Betaproteobacteria* and *Bacteroidetes* were three of the most abundant phyla in freshwater lakes. The high frequency of shared taxa within these groups may be related to an increased probability of dispersal between habitat types with large population sizes. A second commonality across these groups is that they all include photoheterotrophic organisms. Each group includes rhodopsin-containing bacteria and the proteobacterial classes are the primary aerobic anoxygenic phototrophs in aquatic systems (3, 26-30). A recently described novel rhodopsin clade, acidirhodopsin, is found in marine *Acidimicrobiales* and is related to freshwater actinorhodopsins (4). Photoheterotrophic lifestyles may enable microbial cells to persist in an environment until conditions arise that allow population sizes to expand. Notably, *Actinobacteria* has lower sequence similarity between pairs of marine and freshwater samples and contains fewer shared nodes than the other four groups. Below-average growth rates (31) and the dependence of some actinobacterial lineages on other bacteria (32) may contribute to apparent differences in the ability of *Actinobacteria* and other abundant taxa to transition between marine and freshwater systems.

Bacteria from groups with comparatively high transition frequency employ a wide range of aquatic lifestyles, from small streamlined SAR11 cells that harvest low-molecular weight dissolved organic matter (33) to particle-attached *Bacteriodetes* with the ability to degrade polymers and genes for gliding motility (34). Transitions between freshwater and marine systems likely included lineages that developed novel adaptations in one habitat that facilitated success in the other habitat by conferring a degree of selective advantage or niche complementarity. Some lineages that successfully transitioned between marine and freshwater habitats may have been aided by an ability to thrive under a range of salinities. Aquatic bacterial strains identified as salinity generalists by Matias and colleagues (35) include representatives of the Comamonadaceae (*Betaproteobacteria*), Pseudomonadaceae (*Gammaproteobacteria*) and Vibrionaceae (*Gammaproteobacteria*); all three of these families contained shared MED nodes in our meta-analysis (13, 4, and 5 nodes, respectively).

### Insights from the SAR11 clade

Our observation that freshwater lakes contain non-LD12 SAR11 (*Alphaproteobacteria*) with abundances near detection limits challenges the classic understanding of marine-freshwater transitions. SAR11 sequences classified to groups other than the freshwater LD12 group were found in three different lake systems: a humic lake in Northern Wisconsin, a Tibetan Plateau lake, and multiple Laurentian Great Lakes. The same non-LD12 SAR11 node was detected in Lake Michigan and Lake Ontario samples collected from multiple years, depths, and stations within each lake. The persistent observation of this node suggests that there is an established population in the Great Lakes, presumably making ecological contributions. The low relative abundance of non-LD12 SAR11 in the Great Lakes was exacerbated by the 515F-C/806R primer set bias against SAR11 (36). Using 515F/926R primers, we observed an 8-30x increase in the relative abundance non-LD12 SAR11 sequences (Paver et al. in prep.); however these primers have not yet been widely employed, so we relied on datasets using the more common 515F/806R primers for our meta-analysis.

Organisms with abundances at or near the detection limits of current sequencing practices are frequently removed from analyses that exclude sequences below a specified abundance threshold (37). However, populations with low representation in sequencing libraries may have unintuitively large census population sizes in a system. A population with a density of one cell per ml has a population size of billions of cells in a one meter depth layer of a small lake like Trout Bog and quadrillions of cells in a one meter depth layer of Lake Michigan. Assuming that the probability of sequencing is proportional to cell abundance and there are 500,000 cells per ml, a sequence from that population will not be detected 74% of the time and a single sequence will be detected 22% of the time from a sample with 150,000 sequences (slightly higher than any samples included in our meta-analysis), making the population likely to go unreported. Aquatic systems contain a systematically overlooked pool of diversity that may harbor organisms that immigrated from other habitats but have not become dominant in the system. These frequently overlooked low abundance taxa may make disproportionately large contributions to ecosystem function (38) and could serve as a source of taxa available to take advantage of changing environmental conditions, akin to what Shade and colleagues (39) describe as ‘conditionally rare taxa.’

The perception that very few marine-freshwater transitions occurred during the evolutionary diversification of SAR11 was based on observations where all SAR11 detected in freshwater systems were classified to the freshwater LD12 group while no marine sequences were identified as LD12 (40). The first indication that non-LD12, marine-like SAR11 inhabit lakes came from a recently reconstructed parial genome classified as SAR11 subtype I/II from Lake Baikal (41). The partial genome from Lake Baikal combined with our observations provide robust evidence of SAR11 lineages other than LD12 (SAR11 IIIb) inhabiting freshwater lakes. Prior to these findings, marine SAR11 had been observed in saline lakes (40, 42) and LD12 had been detected and determined to assimilate thymidine in the brackish Gulf of Gdańsk, which is located on the southern Baltic Sea coast and receives freshwater input from the Vistula River (43). The first cultured representative of LD12 was isolated from the coastal lagoon of Lake Borgne (44). Within the SAR11 clade, there appears to be a fine-tuning of salinity preference, resulting in lineage-specific distributions along salinity gradients (10, 45).

We were not able to determine the exact placement of most Great Lakes non-LD12 SAR11 within the larger SAR11 clade phylogeny based on the phylogenetic placement of short metagenomic reads. The small fraction of Great Lakes non-LD12 SAR11 sequences that were classified to the clade level were split between clades Ia and IIIa, a sister group of LD12 commonly observed in brackish environments (10, 45, 46). Great Lakes metagenome sequences were not classified to the genome from Lake Baikal, suggesting that the non-LD12 SAR11 populations in Lake Baikal and the Laurentian Great Lakes represent separate transitions. We found a high variability in the fraction of Great Lakes sequences classified as marine SAR11 across protein clusters, which could be explained by the lack of a highly similar reference genome, or by horizontal gene transfer between LD12 and marine SAR11 populations within the Great Lakes. While adaptation to environments with specific salinities appears to have played a major role in determining the distribution of SAR11 organisms, the prior perception that LD12 is the only freshwater Pelagibacterales appears to have been an artifact of undersampling.

As Logares and colleagues (7) discussed, the potential for SAR11 to disperse to freshwater systems is high due to the immense population sizes of marine SAR11 and postulated potential of microbes to disperse long distances. Walsh and colleagues (47) speculated that ancestors of freshwater LD12 may have been successful in transitioning from the ocean to freshwater due to a lesser dependency on Na+ compared to other marine lineages, which may have been a contributing factor for other SAR11 lineages crossing over into freshwater as well. The LD12 lineage originated from marine populations, acquiring genomic adaptations, including loss of specific carbon degradation pathways and changes to central carbon metabolism, that enabled populations to reach high abundances in certain lakes and become widely distributed in lakes across the globe (48, 49). Based on our results, other SAR11 lineages which had been observed exclusively in brackish and higher salinity environments appear to have made the transition to freshwater without dominating ecosystems.

### Using 16S rRNA datasets to detect transitions

There are several important caveats to consider when comparing microbial diversity between habitat types. First, we can only survey abundant, extant diversity. Analyzing 16S rRNA amplicon datasets gave us the important benefit of deep sequencing relative to other sequence-based approaches; however, the 16S rRNA gene is not a good marker for differentiating closely related organisms (50). Marine and freshwater microorganisms classified as ‘shared’ in our analyses likely have habitat-specific differentiation at the genome level. Given the substantial overlap that we observed between marine and freshwater 16S rRNA amplicon sequences, recent and frequent transitions cannot be ruled out on the basis of 16S rRNA datasets. Microbial community composition can also be affected by biases, including those resulting from DNA extraction method (51) and 16S rRNA gene primer set. For example, the 515F-C/806R primer set used by many datasets included in our meta-analysis has documented biases against SAR11 and *Thaumarchaeota* (36, 52). We minimized the effects of these biases on our comparative analyses by focusing on the sequence level and discarding abundance data. Taxa that were not amplified by primer sets used in the datasets analyzed are beyond the scope of this work. Another consideration is the possibility that shared taxa arise due to reagent contamination, for example in DNA extraction kits (53) or cross-contamination from multi-sample multiplexing (54). Given that most samples were extracted with study-specific methods including modifications to commercial kits or no commercial kit and sequenced independently for each study, as well as the repeatability of our results across samples, it is unlikely that our observations of shared taxa can be explained by contamination.

### Summary

Marine and freshwater systems are phylogenetically distinct, while at the same time harboring taxa that appear in both environments. Some taxonomic groups appear to be exclusive to marine or freshwater environments. At the same time, some taxonomic units appear in both habitat types; we identified 171 shared MED nodes across marine and freshwater habitats. It remains to be seen whether individual cells with marine-like or freshwater-like 16S rRNA resemble populations found in the other habitat genome-wide, or whether there is genomic mosaicism. Families at the extremes – lineages with a high degree of habitat-specific diversification or a large number of taxa found in both habitats – may serve as targets for future work investigating ecological plasticity and/or adaptations and the evolution of microbial lineages. There is clearly precedent and potential for marine and freshwater organisms to transition between habitats with different salinities and adapt to new environmental conditions, available resources, and interactions with neighboring organisms. A bank of near- or below-detection diversity, including cross-system immigrant populations, may contribute to community genomic diversity via horizontal gene transfer and exploit opportunities for niche expansion as environmental conditions change.

## Methods

### 16S rRNA sequence processing

We carried out a meta-analysis of marine and freshwater 16S rRNA gene sequencing datasets spanning the V4 region (Table 2; Table S2). For a dataset to be included in our analysis, sequence reads needed to encompass bases 515 through 805 of the 16S rRNA gene. We augmented publically available datasets with samples from the Laurentian Great Lakes sequenced by the Joint Genome Institute (Paver et al., in prep). Sequence processing was carried out using mothur v 1.38.1 unless otherwise noted (55). We merged paired sequence reads using make.contigs and quality filtered single reads using trim.seqs (window size = 50, minimum average quality score = 35). All sequences were then combined and processed following a modified version of the mothur MiSeq standard operating protocol accessed 27 September 2016 (56). Screening retained 200-300 bp sequences with no ambiguities and maximum homopolymer stretches fewer than 24 bases. Screened sequences were aligned to the Silva v128 reference alignment (57, 58) prior to chimera identification using uchime (59) as implemented by mothur. Identified chimeras were subsequently removed. Sequences were classified in mothur using Silva v128, and those identified as ‘Chloroplast’, ‘Mitochondria’, ‘unknown’, or ‘Eukaryota’ were removed from the dataset. To generate an estimate of taxon relatedness independent of operationally selected sequence identity cutoffs, we implemented direct comparisons for all sequences within each phylum and proteobacterial class. Pairwise sequence distances were calculated and furthest-neighbor clustering was used to group sequences into taxonomic units for all sequence identity cutoff values from unique down to the level where all sequences converge into one taxonomic unit at a precision of 1000. For groups with distance matrices too large to process all sequences together (*Alphaproteobacteria*, *Bacteroidetes*, *Betaproteobacteria*, *Gammaproteobacteria*), cluster.split was implemented at the order level (taxonomic level 4) to group sequences into taxonomic units at cutoff values from unique down to 0.30. Classification-based cluster splitting was not a feasible approach for the *Actinobacteria*, so pre.cluster was run on sequences prior to calculating furthest-neighbor clusters. We also implemented Minimum Entropy Decomposition (MED), a method that employs Shannon entropy to partition sequences into taxonomic units referred to as ‘nodes’ using information-rich nucleotide positions and ignoring stochastic variation (37). We ran MED with a minimum substantive abundance of 10 sequences and 4 discriminant locations.

**Table 2.**
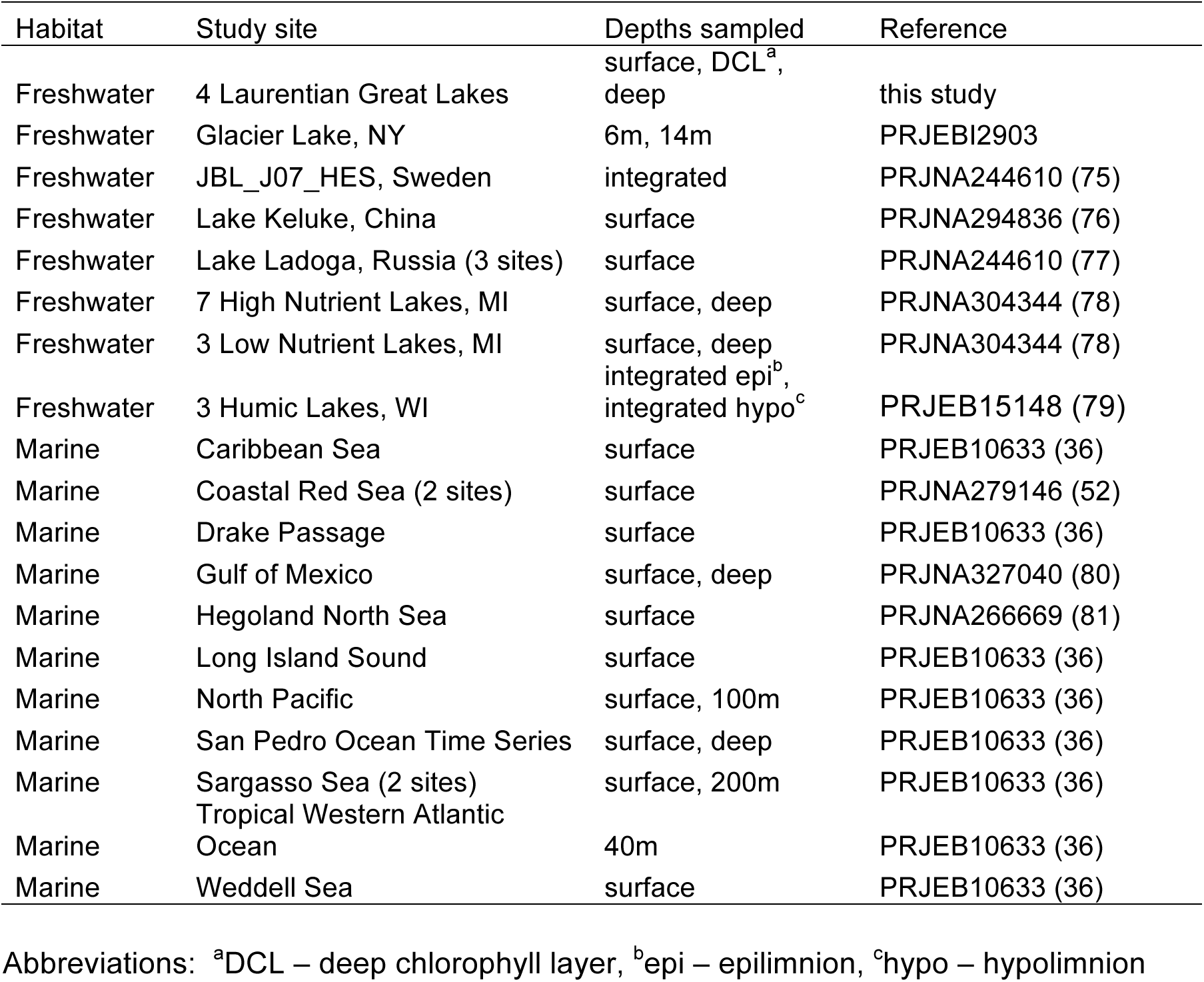
16S rRNA v4 region sequencing datasets included in the meta-analysis

### Statistical and phylogenetic comparisons of marine and freshwater datasets

We compared marine and freshwater samples and sequences at the levels of taxonomic classification, MED nodes, and sequence similarity using R version 3.3.2 (R Core Team 2016). Phyla and proteobacterial classes that were differentially abundant in marine vs. freshwater samples were identified by testing for differences in the log2 fold change using a parametric Wald test implemented by DESeq2 (60).

We generated a maximum-likelihood tree from mothur-aligned sequences using the GTRGAMMA model in RaxML v7.7.9 (61). To visualize sub-trees and calculate UniFrac distances for specific groups, bacterial trees were rooted with a Marine Group I archaeal sequence (A000001667) using the ape R package (62). Trees were visualized using the interactive Tree Of Life (63). Sequences were rarefied and data were subset for subsequent analyses using the phyloseq package (64). We rarefied samples to even depth (9 827 sequences/ sample) and calculated unweighted UniFrac distances for each pair of samples using GUniFrac (65). We tested the significance of UniFrac distances between marine and freshwater samples using Permutational Multivariate Analysis of Variance (PerMANOVA) as implemented by the adonis function in the vegan package (Oksanen et al. 2016). The same approach was used to calculate UniFrac values for each phylum and proteobacterial class that was observed in at least three marine and three freshwater samples and contained at least five MED nodes. Samples containing fewer than 500 sequences for a given phylum or class were removed from the analysis. We combined all sequences from each habitat and calculated unweighted UniFrac distances for phyla, proteobacterial classes, orders and families using GUniFrac following sequence rarefaction to even depth. To test the significance of calculated UniFrac values for each phylum and proteobacterial class, unweighted UniFrac values were calculated for 1000 independent swap randomizations of the presence-absence sample matrix generated by the randomizeMatrix function in the picante package (66). Using these distances as a null distribution, one sample z tests were conducted to test the null hypothesis that the phylogenetic tree is not grouped by habitat. A Bonferroni correction was implemented as a conservative measure to account for multiple testing in determining significance.

We calculated Jaccard distances for pairs of unrarefied samples at each sequence identity cutoff for each phylum and proteobacterial class. The Jaccard distance is calculated as 1 minus the intersection of two samples (i.e., the number of shared units) divided by the union of two samples (i.e., the total number of units); a Jaccard distance of 1 means that no taxa are shared between samples. The sequence identity cutoff value at which marine and freshwater sample pairs first contain shared taxa (Jaccard distance < 1) was summarized for each phylum/ class using boxplots.

We identified ‘shared nodes’ as those observed in the unrarefied sequence set for at least one marine and at least one freshwater sample. For each phylum and proteobacterial class containing more than five nodes observed in both marine and freshwater samples (i.e., shared nodes), we generated accumulation curves to visualize the number of shared MED nodes as a function of the number of freshwater or marine sampling sites using the specaccum function in the vegan package (Oksanen et al. 2016).

### Metagenomic evidence of non-LD12 SAR11 in the Great Lakes

To test whether there is genome-wide evidence beyond the 16S rRNA locus for non-LD12 SAR11 cells in the Great Lakes, we identified reads that mapped with high confidence to non-LD12 SAR11 in a SAR11/LD12 reference phylogeny. Briefly, we analyzed metagenomes from samples collected from the surface of each of the Great Lakes in spring 2012 (IMG database project IDs: Ga0049080-Ga0049085) as well as a marine sample from the Tara Oceans project collected from the North Atlantic Ocean Westerlies Biome near Bermuda for comparison (accession number ERR599123) (67). Sequence reads from each metagenome were searched against the nr protein database (68 downloaded 13 April 2017); using diamond v. 0.8.18.80 (69) and sequences whose best hit matched the *Pelagibacterales* family were identified using Krona Tools v. 2.7 (70) and extracted. These putative *Pelagibacterales* reads were then mapped to a database of SAR11/LD12 protein clusters using a translated query-protein subject (blastx) BLAST v. 2.2.28 (71) with e-value <0.001 and alignment length >60 amino acids (>50 amino acids for the Tara Oceans sample since sequence reads were shorter). Our protein clusters database was constructed from publically available SAR11 and LD12 genomes (Table 3) using all-vs-all blastp and MCL clustering (72) as implemented by anvi’o v. 2.4.0 (73). We focused our analysis on putative core protein clusters found single-copy in at least 6 SAR11 genomes and 3 LD12 genomes; notably all but one LD12 genome derives from single-cell genome amplification and sequencing and are therefore incomplete. For each protein cluster, we backtranslated amino acid alignments, generated by muscle v. 3.8 (74) within anvi’o, to nucleotide alignments. We then used RAxML v. 7.2.6 with the GTRGAMMA model to generate a maximum likelihood tree for each protein cluster based on its corresponding nucleotide alignment (61). Metagenomic reads that mapped to each protein cluster by blastx were aligned to the cluster’s reference nucleotide alignment using HMMER v 3.1b2 (hmmer.org). Reads were then classified taxonomically using pplacer with the -p flag to calculate prior probabilities and guppy classify using the -pp flag to use posterior probability for the pplacer classifier criteria v1.1.alpha19-0-g807f6f3 (22). The reference package for taxonomy classification was generated using taxtastic v 0.5.4 (http://fhcrc.github.io/taxtastic/index.html). We analyzed the resulting databases using the R package BoSSA v2.1 (https://cran.rproject.org/web/packages/BoSSA/index.html).

**Table 3.**
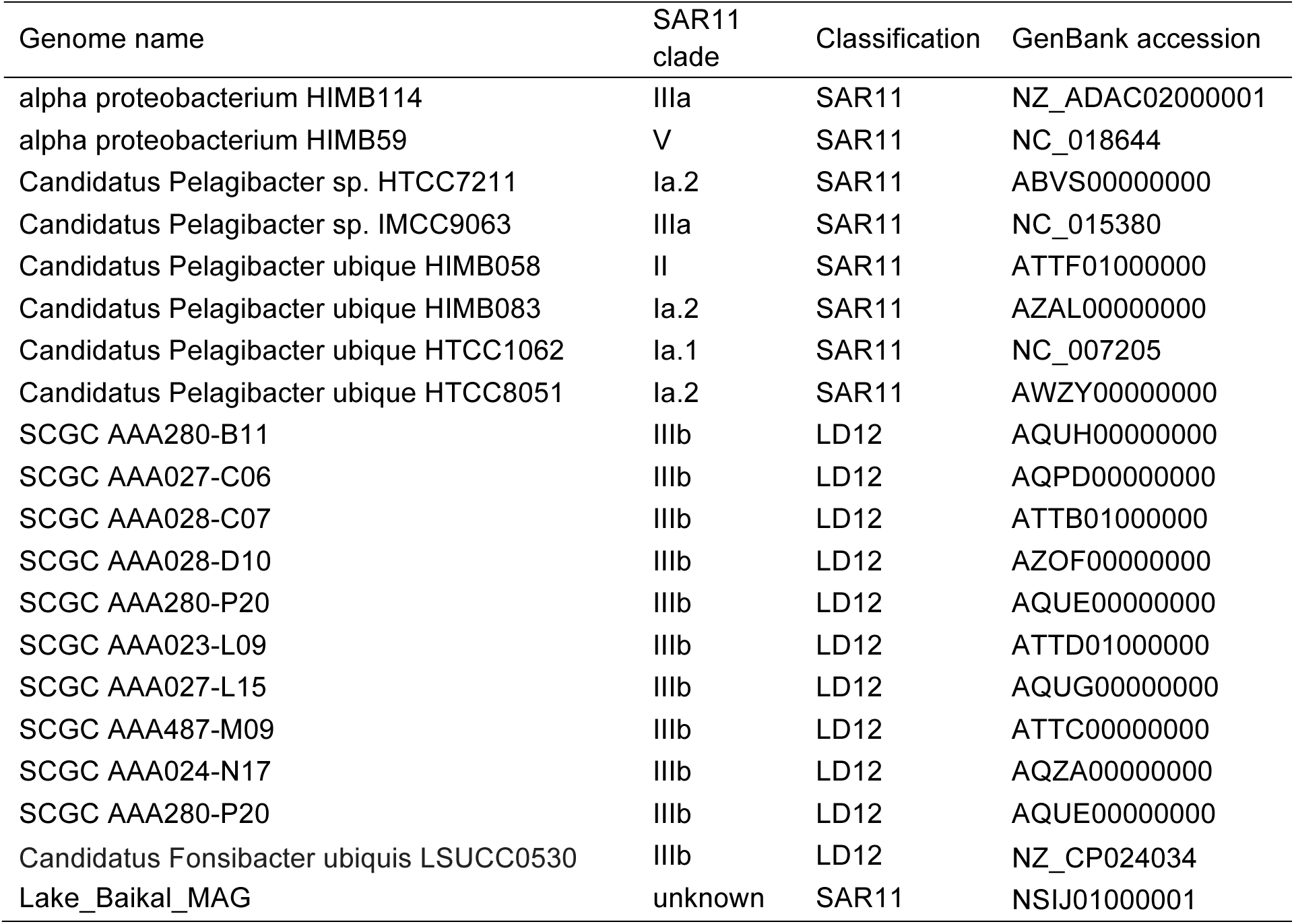
SAR11 and LD12 genomes included in pangenomic analysis.

Data availability: Laurentian Great Lake sequences have been deposited under accession numbers (to be added during revision). R code and associated data files are available at https://bitbucket.org/greatlakes/marinefwmeta.git. Intermediate data files referenced in code, but too large to store on bitbucket will be made available in a public data repository (digital object identifiers to be added during revision) and can be accessed at the following link for review purposes: https://uchicago.box.com/s/o4vjbe1wmf0umj87jw5bhs2w0y6dtdvq

## Acknowledgements

Great Lakes samples were sequenced by the DOE Joint Genome Institute (CSP #1565) and computational resources were provided by the University of Chicago Research Computing Center. We thank Glenn Warren and the science staff in the Great Lakes National Program Office of the US EPA and the Captain and crew of the R/V *Lake Guardian* for facilitating sample collection on the Great Lakes; A. M. Eren for assistance with MED; J. Podowski for Krona Tools output; and Jacob Waldbauer and the Coleman and Waldbauer lab group at UChicago for constructive feedback.

## Author contributions

Study conception and design: SFP, MLC; analysis and interpretation of data: SFP, DJM, RJN, MLC; drafting of manuscript: SFP, MLC; critical revision: SFP, DJM, RJN, MLC.

## Supplementary Tables

Table S1. UniFrac distance calculated between marine and freshwater sequences for each family.

Table S2. Additional information on 16S rRNA tag sequencing datasets compiled for the meta-analysis.

## Supplementary Figures

Figure S1. Principal coordinate analysis of unweighted UniFrac distances between marine and freshwater assemblages characterized by 16S rRNA V4 gene sequences.

Figure S2. Median relative abundance of abundant orders orders (a) and families (b) in surface waters of freshwater datasets compared to marine datasets (bars indicate ranges).

Figure S3. Median unweighted UniFrac distances (bars indicate range) between pairs of freshwater and marine samples compared to unweighted UniFrac distance between combined freshwater and combined marine samples for each phylum and proteobacterial class (a). Unweighted UniFrac distances between combined freshwater and combined marine samples for orders (b) and families (c) within each phylum/ class. Phyla, classes, orders, and families needed to be present in at least 3 freshwater and 3 marine samples and contain 5 nodes.

Figure S4. As sequence identity cutoff value increases, more taxa are shared between marine and freshwater sample pairs. Decrease in Jaccard index between marine and freshwater sample pairs (unshared taxa)/(shared taxa) with increasing sequence similarity cutoff values for *Alphaproteobacteria* and *Verrucomicrobia* (a). Example phylogenetic trees are shown for groups with high sequence similarity (b) and low sequence similarity (c) between marine (blue) and freshwater (green) samples.

Figure S5. Minimum sequence identity threshold (i.e., finest scale resolution) at which pairs of marine and freshwater samples share common taxa. In contrast to Figure 2, this analysis used a dataset where sequences were pre-clustered to allow up to 2 base pair differences within a sequence group. Box plots indicate the median, quartiles, and range of values observed for all marine and freshwater sample pairs. A heatmap illustrates the number of freshwater (F) and marine (M) samples containing representatives of each phyla/ proteobacterial class.

Figure S6. Phylogenetic trees of select phyla and proteobacterial classes: (a) *Alphaproteobacteria*, (b) *Actinobacteria*, (c) *Betaproteobacteria*, (d) *Gammaproteobacteria*, (e) *Cyanobacteria*, (f) *Chloroflexi*. External ring colors indicath the habitats where MED nodes were detected in the dataset: freshwater (green), marine (blue), both (orange). Notable clades are indicated by colored wedges.

Figure S7. Fraction of freshwater nodes shared with marine systems as a function of freshwater sites sampled (a) and fraction of marine nodes shared with freshwater systems as a function of marine sites sampled (b).

Figure S8. Neighbor-joining consensus tree of nucleotide sequences (a) and nucleotide and protein alignment (b) of partial genes from the protein cluster identified as COG2609 (Pyruvate dehydrogenase complex, dehydrogenase E1 component). Strain names are colored based on phylogenetic classification within the SAR11 clade: green - LD12 sequences from group IIIb; light blue - group IIIa, sister group to IIIb; blue - all other marine SAR11 clades included in the analysis (Ia, II); black - a sequence read from the Lake Erie metagenome identified by pplacer as marine SAR11. Consensus support values (%) are indicated on tree branches (a). Vertical boxes have been added to the alignment to indicate amino acids where the Great Lakes sequence contains shared characters with a subset of included strains.

## References

1. Grossart H-P. 2010. Ecological consequences of bacterioplankton lifestyles: changes in concepts are needed. Environmental Microbiology Reports 2:706–714.

2. Cole JJ. 1982. Interactions between bacteria and algae in aquatic ecosystems. Annual Review of Ecology and Systematics 13:291–314.

3. Koblížek M. 2015. Ecology of aerobic anoxygenic phototrophs in aquatic environments. FEMS Microbiology Reviews 39:854–870.

4. Mizuno CM, Rodriguez-Valera F, Ghai R. 2015. Genomes of Planktonic Acidimicrobiales: Widening Horizons for Marine Actinobacteriaby Metagenomics. mBio 6:e02083–14–11.

5. Salcher MM, Neuenschwander SM, Posch T, Pernthaler J. 2015. The ecology of pelagic freshwater methylotrophs assessed by a high-resolution monitoring and isolation campaign. The ISME Journal 9:2442–2453.

6. Martiny JBH, Jones SE, Lennon JT, Martiny AC. 2015. Microbiomes in light of traits: A phylogenetic perspective. Science 350:aac9323–aac9323.

7. Logares R, Bråte J, Bertilsson S, Clasen JL, Shalchian-Tabrizi K, Rengefors K. 2009. Infrequent marine–freshwater transitions in the microbial world. Trends in Microbiology 17:414–422.

8. Lozupone CA, Knight R. 2007. Global patterns in bacterial diversity. Proc Natl Acad Sci USA 104:11436–11440.

9. Thompson LR, Sanders JG, McDonald D, Amir A, Ladau J, Locey KJ, Prill RJ, Tripathi A, Gibbons SM, Ackermann G, Navas-Molina JA, Janssen S, Kopylova E, Vázquez-Baeza Y, Gonzalez A, Morton JT, Mirarab S, Zech Xu Z, Jiang L, Haroon MF, Kanbar J, Zhu Q, Jin Song S, Kosciolek T, Bokulich NA, Lefler J, Brislawn CJ, Humphrey G, Owens SM, Hampton-Marcell J, Berg-Lyons D, McKenzie V, Fierer N, Fuhrman JA, Clauset A, Stevens RL, Shade A, Pollard KS, Goodwin KD, Jansson JK, Gilbert JA, Knight R, Rivera JLA, Al-Moosawi L, Alverdy J, Amato KR, Andras J, Angenent LT, Antonopoulos DA, Apprill A, Armitage D, Ballantine K, Bárta J, Baum JK, Berry A, Bhatnagar A, Bhatnagar M, Biddle JF, Bittner L, Boldgiv B, Bottos E, Boyer DM, Braun J, Brazelton W, Brearley FQ, Campbell AH, Caporaso JG, Cardona C, Carroll J, Cary SC, Casper BB, Charles TC, Chu H, Claar DC, Clark RG, Clayton JB, Clemente JC, Cochran A, Coleman ML, Collins G, Colwell RR, Contreras M, Crary BB, Creer S, Cristol DA, Crump BC, Cui D, Daly SE, Davalos L, Dawson RD, Defazio J, Delsuc F, Dionisi HM, Dominguez-Bello MG, Dowell R, Dubinsky EA, Dunn PO, Ercolini D, Espinoza RE, Ezenwa V, Fenner N, Findlay HS, Fleming ID, Fogliano V, Forsman A, Freeman C, Friedman ES, Galindo G, Garcia L, Garcia-Amado MA, Garshelis D, Gasser RB, Gerdts G, Gibson MK, Gifford I, Gill RT, Giray T, Gittel A, Golyshin P, Gong D, Grossart H-P, Guyton K, Haig S-J, Hale V, Hall RS, Hallam SJ, Handley KM, Hasan NA, Haydon SR, Hickman JE, Hidalgo G, Hofmockel KS, Hooker J, Hulth S, Hultman J, Hyde E, Ibáñez-Álamo JD, Jastrow JD, Jex AR, Johnson LS, Johnston ER, Joseph S, Jurburg SD, Jurelevicius D, Karlsson A, Karlsson R, Kauppinen S, Kellogg CTE, Kennedy SJ, Kerkhof LJ, King GM, Kling GW, Koehler AV, Krezalek M, Kueneman J, Lamendella R, Landon EM, Lane-deGraaf K, LaRoche J, Larsen P, Laverock B, Lax S, Lentino M, Levin II, Liancourt P, Liang W, Linz AM, Lipson DA, Liu Y, Lladser ME, Lozada M, Spirito CM, MacCormack WP, MacRae-Crerar A, Magris M, Martín-Platero AM, Martín-Vivaldi M, Martínez LM, Martínez-Bueno M, Marzinelli EM, Mason OU, Mayer GD, McDevitt-Irwin JM, McDonald JE, McGuire KL, McMahon KD, McMinds R, Medina M, Mendelson JR, Metcalf JL, Meyer F, Michelangeli F, Miller K, Mills DA, Minich J, Mocali S, Moitinho-Silva L, Moore A, Morgan-Kiss RM, Munroe P, Myrold D, Neufeld JD, Ni Y, Nicol GW, Nielsen S, Nissimov JI, Niu K, Nolan MJ, Noyce K, O’Brien SL, Okamoto N, Orlando L, Castellano YO, Osuolale O, Oswald W, Parnell J, Peralta-Sánchez JM, Petraitis P, Pfister C, Pilon-Smits E, Piombino P, Pointing SB, Pollock FJ, Potter C, Prithiviraj B, Quince C, Rani A, Ranjan R, Rao S, Rees AP, Richardson M, Riebesell U, Robinson C, Rockne KJ, Rodriguezl SM, Rohwer F, Roundstone W, Safran RJ, Sangwan N, Sanz V, Schrenk M, Schrenzel MD, Scott NM, Seger RL, Seguin-Orlando A, Seldin L, Seyler LM, Shakhsheer B, Sheets GM, Shen C, Shi Y, Shin H, Shogan BD, Shutler D, Siegel J, Simmons S, Sjöling S, Smith DP, Soler JJ, Sperling M, Steinberg PD, Stephens B, Stevens MA, Taghavi S, Tai V, Tait K, Tan CL, Tas N, Taylor DL, Thomas T, Timling I, Turner BL, Urich T, Ursell LK, van der Lelie D, Van Treuren W, van Zwieten L, Vargas-Robles D, Thurber RV, Vitaglione P, Walker DA, Walters WA, Wang S, Wang T, Weaver T, Webster NS, Wehrle B, Weisenhorn P, Weiss S, Werner JJ, West K, Whitehead A, Whitehead SR, Whittingham LA, Willerslev E, Williams AE, Wood SA, Woodhams DC, Yang Y, Zaneveld J, Zarraonaindia I, Zhang Q, Zhao H. 2017. A communal catalogue reveals Earth’s multiscale microbial diversity. Nature 104:11436–24.

10. Herlemann DP, Labrenz M, Jürgens K, Bertilsson S, Waniek JJ, Andersson AF. 2011. Transitions in bacterial communities along the 2000km salinity gradient of the Baltic Sea. The ISME Journal 5:1571–1579.

11. Fortunato CS, Crump BC. 2015. Microbial Gene Abundance and Expression Patterns across a River to Ocean Salinity Gradient. PLoS ONE 10:e0140578–22.

12. Logares R, m ESLO, Langenheder S, Logue JURB, Paterson H, Laybourn-Parry J, Rengefors K, Tranvik L, Bertilsson S. 2012. Biogeography of bacterial communities exposed to progressive long-term environmental change. The ISME Journal 7:937–948.

13. Zwart G, Crump BC, Agterveld MPK-V, Hagen F, Han S-K. 2002. Typical freshwater bacteria: an analysis of available 16S rRNA gene sequences from plankton of lakes and rivers. Aquat Microb Ecol 28:141–155.

14. Newton RJ, Jones SE, Eiler A, McMahon KD, Bertilsson S. 2011. A guide to the natural history of freshwater lake bacteria. Microbiol Mol Biol Rev 75:14–49.

15. Hubbell SP. 2001. The Unified Neutral Theory of Biodiversity and Biogeography. Princeton University Press.

16. Lenski RE, Rose MR, Simpson SC, Tadler SC. 1991. Long-Term Experimental Evolution in Escherichia coli. I. Adaptation and Divergence During 2,000 Generations. The American Naturalist 138:1315–1341.

17. Elena SF, Lenski RE. 2003. Microbial genetics: Evolution experiments with microorganisms: the dynamics and genetic bases of adaptation. Nat Rev Genet 4:457–469.

18. Janssen J, Marsden JE, Hrabik TR, Stockwell JD. 2014. Are the Laurentian Great Lakes great enough for Hjort? ICES Journal of Marine Science 71:2242–2251.

19. Hecky RE, Campbell P, Hendzel LL. 1993. The stoichiometry of carbon, nitrogen, and phosphorus in particulate matter of lakes and oceans. Limnol Oceanogr 38:709–724.

20. Paytan A, McLaughlin K. 2007. The Oceanic Phosphorus Cycle. Chem Rev 107:563–576.

21. Sterner RW. 2010. In situ-measured primary production in Lake Superior. Journal of Great Lakes Research 36:139–149.

22. Matsen FA, Kodner RB, Armbrust EV. 2010. pplacer: linear time maximum-likelihood and Bayesian phylogenetic placement of sequences onto a fixed reference tree. BMC Bioinformatics 11:538.

23. Eiler A, Zaremba-Niedzwiedzka K, Martínez-García M, McMahon KD, Stepanauskas R, Andersson SGE, Bertilsson S. 2013. Productivity and salinity structuring of the microplankton revealed by comparative freshwater metagenomics. Environ Microbiol 16:2682–2698.

24. Barberán A, Casamayor EO. 2010. Global phylogenetic community structure and β-diversity patterns in surface bacterioplankton metacommunities. Aquat Microb Ecol 59:1–10.

25. Lozupone C, Knight R. 2005. UniFrac: a new phylogenetic method for comparing microbial communities. Applied and Environmental Microbiology 71:8228–8235.

26. Gómez-Consarnau L, Akram N, Lindell K, Pedersen A, Neutze R, Milton DL, González JM, Pinhassi J. 2010. Proteorhodopsin Phototrophy Promotes Survival of Marine Bacteria during Starvation. PLoS Biol 8:e1000358–10.

27. Brindefalk B, Ekman M, Ininbergs K, Dupont CL, Yooseph S, Pinhassi J, Bergman B. 2016. Distribution and expression of microbial rhodopsins in the Baltic Sea and adjacent waters. Environ Microbiol 18:4442–4455.

28. Martínez-García M, Swan BK, Poulton NJ, Gomez ML, Masland D, Sieracki ME, Stepanauskas R. 2011. High-throughput single-cell sequencing identifies photoheterotrophs and chemoautotrophs in freshwater bacterioplankton. The ISME Journal 6:113–123.

29. Béjà O, Suzuki MT, Heidelberg JF, Nelson WC, Preston CM, Hamada T, Eisen JA, Fraser CM, DeLong EF. 2002. Unsuspected diversity among marine aerobic anoxygenic phototrophs. Nature 415:630–633.

30. Yutin N, Suzuki MT, Teeling H, Weber M, Venter JC, Rusch DB, Béjà O. 2007. Assessing diversity and biogeography of aerobic anoxygenic phototrophic bacteria in surface waters of the Atlantic and Pacific Oceans using the Global Ocean Sampling expedition metagenomes. Environ Microbiol 9:1464–1475.

31. Šimek K, Horňak K, Jezbera J, Nedoma J, Vrba J, Straškrabova V, Macek M, Dolan JR, Hahn MW. 2006. Maximum growth rates and possible life strategies of different bacterioplankton groups in relation to phosphorus availability in a freshwater reservoir. Environ Microbiol 8:1613–1624.

32. Hahn MW. 2009. Description of seven candidate species affiliated with the phylum Actinobacteria, representing planktonic freshwater bacteria. Int J Syst Evol Microbiol 59:112–117.

33. Giovannoni SJ. 2017. SAR11 Bacteria: The Most Abundant Plankton in the Oceans. Annu Rev Mar Sci 9:231–255.

34. Fernández-Gómez B, Richter M, Schüler M, Pinhassi J, Acinas SG, González JM, Pedrós-Alió C. 2013. Ecology of marine Bacteroidetes: a comparative genomics approach. The ISME Journal 7:1026–1037.

35. Matias MG, Combe M, Barbera C, Mouquet N. 2012. Ecological strategies shape the insurance potential of biodiversity. Front Microbiol 3:432.

36. Parada AE, Needham DM, Fuhrman JA. 2016. Every base matters: assessing small subunit rRNA primers for marine microbiomes with mock communities, time series and global field samples. Environ Microbiol 18:1403–1414.

37. Eren AM, Morrison HG, Lescault PJ, Reveillaud J, Vineis JH, Sogin ML. 2015. Minimum entropy decomposition: unsupervised oligotyping for sensitive partitioning of high-throughput marker gene sequences. The ISME Journal 9:968–979.

38. Jousset A, Bienhold C, Chatzinotas A, Gallien L, Gobet A, Kurm V, Küsel K, Rillig MC, Rivett DW, Salles JF, van der Heijden MGA, Youssef NH, Zhang X, Wei Z, Hol WHG. 2017. Where less may be more: how the rare biosphere pulls ecosystems strings. The ISME Journal 1–10.

39. Shade A, Jones SE, Caporaso JG, Handelsman J, Knight R, Fierer N, Gilbert JA. 2014. Conditionally rare taxa disproportionately contribute to temporal changes in microbial diversity. mBio 5:e01371–14.

40. Logares R, Bråte J, Heinrich F, Shalchian-Tabrizi K, Bertilsson S. 2010. Infrequent transitions between saline and fresh waters in one of the most abundant microbial lineages (SAR11). Molecular Biology and Evolution 27:347–357.

41. Cabello-Yeves PJ, Zemskaya TI, Rosselli R, Coutinho FH, Zakharenko AS, Blinov VV, Rodriguez-Valera F. 2018. Genomes of Novel Microbial Lineages Assembled from the Sub-Ice Waters of Lake Baikal. Applied and Environmental Microbiology 84:e02132–17–21.

42. Oh S, Zhang R, Wu QL, Liu W-T. 2016. Evolution and adaptation of SAR11 and Cyanobiumin a saline Tibetan lake. Environmental Microbiology Reports 1–26.

43. Piwosz K, Salcher MM, Zeder M, Ameryk A, Pernthaler J. 2013. Seasonal dynamics and activity of typical freshwater bacteria in brackish waters of the Gulf of Gdansk. Limnol Oceanogr 58:817–826.

44. Henson MW, Lanclos VC, Faircloth BC, Thrash JC. 2018. Cultivation and genomics of the first freshwater SAR11 (LD12) isolate. The ISME Journal 420:806.

45. Ortmann AC, Santos TTL. 2016. Spatial and temporal patterns in the Pelagibacteraceae across an estuarine gradient. FEMS Microbiol Ecol 92:fiw133.

46. Kan J, Evans SE, Chen F, Suzuki MT. 2008. Novel estuarine bacterioplankton in rRNA operon libraries from the Chesapeake Bay. Aquat Microb Ecol 51:55–66.

47. Walsh DA, Lafontaine J, Grossart H-P. 2013. On the Eco-Evolutionary Relationships of Fresh and Salt Water Bacteria and the Role of Gene Transfer in Their Adaptation, pp. 55–77. *In* On the eco-evolutionary relationships of fresh and salt water bacteria and the role of gene transfer in their adaptation. Springer New York, New York, NY.

48. Zaremba-Niedzwiedzka K, Viklund J, Zhao W, Ast J, Sczyrba A, Woyke T, McMahon K, Bertilsson S, Stepanauskas R, Andersson SGE. 2013. Single-cell genomics reveal low recombination frequencies in freshwater bacteria of the SAR11 clade. Genome Biology 14:R130.

49. Eiler A, Mondav R, Sinclair L, Fernandez-Vidal L, Scofield DG, Schwientek P, Martínez-García M, Torrents D, McMahon KD, Andersson SG, Stepanauskas R, Woyke T, Bertilsson S. 2016. Tuning fresh: radiation through rewiring of central metabolism in streamlined bacteria. The ISME Journal 1–13.

50. Lan Y, Rosen G, Hershberg R. 2016. Marker genes that are less conserved in their sequences are useful for predicting genome-wide similarity levels between closely related prokaryotic strains. Microbiome 1–13.

51. Vishnivetskaya TA, Layton AC, Lau MCY, Chauhan A, Cheng KR, Meyers AJ, Murphy JR, Rogers AW, Saarunya GS, Williams DE, Pfiffner SM, Biggerstaff JP, Stackhouse BT, Phelps TJ, Whyte L, Sayler GS, Onstott TC. 2013. Commercial DNA extraction kits impact observed microbial community composition in permafrost samples. FEMS Microbiol Ecol 87:217–230.

52. Apprill A, McNally S, Parsons R, Weber L. 2015. Minor revision to V4 region SSU rRNA 806R gene primer greatly increases detection of SAR11 bacterioplankton. Aquat Microb Ecol 75:129–137.

53. Salter SJ, Cox MJ, Turek EM, Calus ST, Cookson WO, Moffatt MF, Turner P, Parkhill J, Loman NJ, Walker AW. 2014. Reagent and laboratory contamination can critically impact sequence-based microbiome analyses. BMC Biology 12:118–12.

54. Wright ES, Vetsigian KH. 2016. Quality filtering of Illumina index reads mitigates sample cross-talk. BMC Genomics 1–7.

55. Schloss PD, Westcott SL, Ryabin T, Hall JR, Hartmann M, Hollister EB, Lesniewski RA, Oakley BB, Parks DH, Robinson CJ, Sahl JW, Stres B, Thallinger GG, Van Horn DJ, Weber CF. 2009. Introducing mothur: open-source, platform-independent, community-supported software for describing and comparing microbial communities. Applied and Environmental Microbiology 75:7537–7541.

56. Kozich JJ, Westcott SL, Baxter NT, Highlander SK, Schloss PD. 2013. Development of a dual-index sequencing strategy and curation pipeline for analyzing amplicon sequence data on the MiSeq Illumina sequencing platform. Applied and Environmental Microbiology 79:5112–5120.

57. Quast C, Pruesse E, Yilmaz P, Gerken J, Schweer T, Yarza P, Peplies J, Glockner FO. 2012. The SILVA ribosomal RNA gene database project: improved data processing and web-based tools. Nucleic Acids Res 41:D590–D596.

58. Yilmaz P, Parfrey LW, Yarza P, Gerken J, Pruesse E, Quast C, Schweer T, Peplies J, Ludwig W, Glöckner FO. 2013. The SILVA and “All-species Living Tree Project (LTP)” taxonomic frameworks. Nucleic Acids Res 42:D643–D648.

59. Edgar RC, Haas BJ, Clemente JC, Quince C, Knight R. 2011. UCHIME improves sensitivity and speed of chimera detection. Bioinformatics 27:2194–2200.

60. Love MI, Huber W, Anders S. 2014. Moderated estimation of fold change and dispersion for RNA-seq data with DESeq2. Genome Biology 15:31–21.

61. Stamatakis A. 2014. RAxML version 8: a tool for phylogenetic analysis and post-analysis of large phylogenies. Bioinformatics 30:1312–1313.

62. Paradis E, Claude J, Strimmer K. 2004. APE: Analyses of Phylogenetics and Evolution in R language. Bioinformatics 20:289–290.

63. Letunic I, Bork P. 2016. Interactive tree of life (iTOL) v3: an online tool for the display and annotation of phylogenetic and other trees. Nucleic Acids Res 44:W242–W245.

64. McMurdie PJ, Holmes S. 2013. phyloseq: An R Package for Reproducible Interactive Analysis and Graphics of Microbiome Census Data. PLoS ONE 8:e61217–11.

65. Chen J, Bittinger K, Charlson ES, Hoffmann C, Lewis J, Wu GD, Collman RG, Bushman FD, Li H. 2012. Associating microbiome composition with environmental covariates using generalized UniFrac distances. Bioinformatics 28:2106–2113.

66. Kembel SW, Cowan PD, Helmus MR, Cornwell WK, Morlon H, Ackerly DD, Blomberg SP, Webb CO. 2010. Picante: R tools for integrating phylogenies and ecology. Bioinformatics 26:1463–1464.

67. Sunagawa S, Coelho LP, Chaffron S, Kultima JR, Labadie K, Salazar G, Djahanschiri B, Zeller G, Mende DR, Alberti A, Cornejo-Castillo FM, Costea PI, Cruaud C, d’Ovidio F, Engelen S, Ferrera I, Gasol JM, Guidi L, Hildebrand F, Kokoszka F, Lepoivre C, Lima-Mendez G, Poulain J, Poulos BT, Royo-Llonch M, Sarmento H, Vieira-Silva S, Dimier C, Picheral M, Searson S, Kandels-Lewis S, Tara Oceans coordinators, Bowler C, de Vargas C, Gorsky G, Grimsley N, Hingamp P, Iudicone D, Jaillon O, Not F, Ogata H, Pesant S, Speich S, Stemmann L, Sullivan MB, Weissenbach J, Wincker P, Karsenti E, Raes J, Acinas SG, Bork P. 2015. Ocean plankton. Structure and function of the global ocean microbiome. Science 348:1261359–1261359.

68. Benson DA. 2004. GenBank. Nucleic Acids Res 33:D34–D38.

69. Buchfink B, Xie C, Huson DH. 2014. Fast and sensitive protein alignment using DIAMOND. Nat Meth 12:59–60.

70. Ondov BD, Bergman NH, Phillippy AM. 2011. Interactive metagenomic visualization in a Web browser. BMC Bioinformatics 12:385.

71. Camacho C, Coulouris G, Avagyan V, Ma N, Papadopoulos J, Bealer K, Madden TL. 2009. BLAST+: architecture and applications. BMC Bioinformatics 10:421–9.

72. van Dongen S, Abreu-Goodger C. 2012. Using MCL to Extract Clusters from Networks, pp. 281–295. *In* van Helden, J, Toussaint, A, Thieffry, D (eds.), Using MCL to extract clusters from networks. Springer New York, New York, NY.

73. Eren AM, Esen ÖC, Quince C, Vineis JH, Morrison HG. 2015. Anvi“o: an advanced analysis and visualization platform for ”omics data. PeerJ 3:e1319.

74. Edgar RC. 2004. MUSCLE: multiple sequence alignment with high accuracy and high throughput. Nucleic Acids Res 32:1792–1797.

75. Logue JB, Langenheder S, Andersson AF, Bertilsson S, Drakare S, Lanzén A, Lindström ES. 2012. Freshwater bacterioplankton richness in oligotrophic lakes depends on nutrient availability rather than on species-area relationships. The ISME Journal 6:1127–1136.

76. Zhong Z-P, Liu Y, Miao L-L, Wang F, Chu L-M, Wang J-L, Liu Z-P. 2016. Prokaryotic Community Structure Driven by Salinity and Ionic Concentrations in Plateau Lakes of the Tibetan Plateau. Applied and Environmental Microbiology 82:1846–1858.

77. Skopina M, Pershina E, Andronov E, Vasileva A, Averina S, Gavrilova O, Ivanikova N, Pinevich A. 2015. Diversity of Lake Ladoga (Russia) bacterial plankton inferred from 16S rRNA gene pyrosequencing: An emphasis on picocyanobacteria. Journal of Great Lakes Research 41:180–191.

78. Schmidt ML, White JD, Denef VJ. 2016. Phylogenetic conservation of freshwater lake habitat preference varies between abundant bacterioplankton phyla. Environ Microbiol 18:1212–1226.

79. Linz AM, Crary BC, Shade A, Owens S, Gilbert JA, Knight R, McMahon KD. 2017. Bacterial Community Composition and Dynamics Spanning Five Years in Freshwater Bog Lakes. mSphere 2:e00169–17–14.

80. Mason OU, Canter EJ, Gillies LE, Paisie TK, Roberts BJ. 2016. Mississippi River Plume Enriches Microbial Diversity in the Northern Gulf of Mexico. Front Microbiol 7:115–13.

81. Teeling H, Fuchs BM, Bennke CM, Krüger K, Chafee M, Kappelmann L, Reintjes G, Waldmann J, Quast C, Glöckner FO, Lucas J, Wichels A, Gerdts G, Wiltshire KH, Amann RI. 2016. Recurring patterns in bacterioplankton dynamics during coastal spring algae blooms. Elife 5:e11888.

